# Additional effects of bariatric surgery and metformin on glucose regulation in non-obese insulin-deficient diabetic rats

**DOI:** 10.64898/2026.04.23.720381

**Authors:** Mélanie Hirlemann, Mélanie Garmon, Lara Ribeiro-Parenti, Danielle Bailbé, Alexandra Willemetz, Hounayda El Jindi Shahrour, Jamileh Movassat, Claire Carette, Maude Le Gall

**Author notes:** Correspondence: Maude LE GALL, Inserm UMRS 1149, Université Paris Cité, 75890 Paris Cedex 18, France.

## Abstract

This study investigates the individual and combined effects of Roux-en-Y gastric bypass (RYGB), sleeve gastrectomy (SG), and metformin on glucose regulation in a non-obese, insulin-deficient model of type 2 diabetes.

Female Goto-Kakizaki (GK) rats underwent RYGB, SG, or sham surgery. Three weeks postoperatively, animals received metformin (50 mg/kg/day, 5 days/week) or vehicle for three additional weeks. Glucose tolerance was assessed using a standardized meal test, and insulin sensitivity was evaluated by insulin tolerance test. Plasma levels of GLP-1, GIP, insulin, and leptin were measured.

RYGB and SG reduced body weight, food intake, and leptin levels, and improved fasting glucose, glucose tolerance, insulin sensitivity, and postprandial incretin and insulin secretion. Metformin alone improved glucose tolerance and insulin sensitivity independently of incretin or insulin changes. When combined with surgery, metformin further reduced postprandial glycemic excursions and advanced the glycemic peak but did not enhance insulin sensitivity or hormone secretion beyond surgery alone.

In conclusion, bariatric surgery and metformin independently improve glucose regulation in non-obese diabetic GK rats. Their combination provides additional benefits on postprandial glucose control, despite no additive effects on insulin sensitivity or hormone levels. These findings support the use of metformin as an adjunct to bariatric surgery in insulin-deficient diabetes and highlight the need for longer-term, sex-inclusive studies to enhance translational relevance.

**NEW & NOTEWORTHY:** Bariatric surgery and metformin each improved glucose regulation in non-obese, insulin-deficient female GK rats. Their combination yielded an additional reduction in postprandial glycemic excursions without further enhancing insulin sensitivity or incretin/insulin secretion. These findings reveal that postprandial glucose dynamics can be modulated independently of hormonal or insulin-sensitivity pathways, highlighting distinct and dissociable mechanisms governing glucose homeostasis in an insulin-deficient model.

## INTRODUCTION

Type 2 diabetes (T2D) is a multifactorial disease frequently associated with obesity, yet it also affects individuals with normal or low body weight. Recent data-driven cluster analyses have identified distinct subtypes of T2D, enabling more refined patient stratification and tailored therapeutic strategies (1). Among these, the severe insulin-deficient diabetes (SIDD) cluster is GADA-negative but shares features with autoimmune diabetes, including younger age at onset, relatively low body mass index (BMI), impaired insulin secretion (low HOMA2-B index), and poor glycemic control. Patients in this cluster in the Scandinavian cohorts exhibit the highest rate of metformin use and the shortest time to initiation of second-line therapy. Notably the ADOPT trial revealed that HbA1c trajectories varied across subgroups, with the SIDD patients showing the most sustained benefit from metformin compared to thiazolidinediones (2) (3). This paradox— persistent metformin uses despite profound β-cell dysfunction—warrants further investigation into its mechanisms and efficacy in this context.

Bariatric-metabolic surgery (BMS), including Roux-en-Y gastric bypass (RYGB) and sleeve gastrectomy (SG), has emerged as a powerful intervention for improving glycemic control in T2D when pharmacological treatments fail (4). Current guidelines even recommend considering metabolic surgery in patients with T2D from a BMI >30 kg/m^2^ to improve glycemic outcomes (5). The metabolic benefits of bariatric surgery are attributed to multiple mechanisms, including enhanced secretion of incretin hormones such as glucagon-like peptide 1 (GLP-1) (6) and glucose-dependent insulinotropic peptide (GIP) (7), which stimulate insulin release in response to food intake (8). However, the efficacy of BMS varies across T2D subtypes, with SIDD patients showing lower rates of diabetes remission and a higher likelihood of requiring continued pharmacotherapy (9). Raverdy et al. reported that despite marked HbA1c reduction after surgery, only 27% of SIDD patients achieved remission at one year. This highlights the need to explore adjunctive treatments that could enhance or prolong the metabolic benefits of surgery in this subgroup. Yet, the combined effects of metformin and bariatric surgery have been insufficiently studied both clinically and experimentally (10).

The Goto-Kakizaki (GK) rat, a non-obese, spontaneous model of T2D recapitulates key features of the SIDD phenotype, including profound of β-cell dysfunction and impaired glucose tolerance (11, 12). Although most preclinical studies have traditionally used male animals, sex-specific differences in metabolic responses to surgery and antidiabetics are increasingly recognized (13, 14). Given that nearly 80% of bariatric procedures are performed in female patients, it is particularly relevant to focus on this population. Investigating these interventions in female GK rats therefore provides novel insights and addresses a critical gap in the preclinical literature (15).

In this study, we investigated the effects of RYGB and SG, alone or in combination with metformin, on glucose homeostasis in female GK rats. Our objective was to determine whether metformin could enhance the metabolic outcomes of bariatric surgery, particularly in the context of postprandial glucose regulation, and to assess the potential for additive or synergistic effects in a non-obese and insulin-deficient model.

## MATERIALS AND METHODS

### Animals

All animal studies were conducted in compliance with European Community guidelines and EU directives regarding animal experimentation. The protocol received approval from the local ethics committee and the French Ministry of Higher Education, Research, and Innovation (MESRI APAFIS #34483).

Female Goto-Kakizaki (GK) rats (BFA, Unité B2PE, Paris, France) (11), aged 5 months and weighing 250–300 g, were housed under a 12-hour light/dark cycle with *ad libitum* access to standard chow (Altromin 1324, Genestil, France).

### Study design

Rats were randomly assigned to Roux-en-Y gastric bypass (RYGB, n=33), sleeve gastrectomy (SG, n=27), or sham surgery (n=33) (Fig.1). Three weeks after surgery, each group was subdivided into metformin-treated or control subgroups: RYGB (n=17), SG (n=15), and sham (n=16); RYGB+Met (n=16), SG+Met (n=12), and sham+Met (n=17).

**Figure 1.**
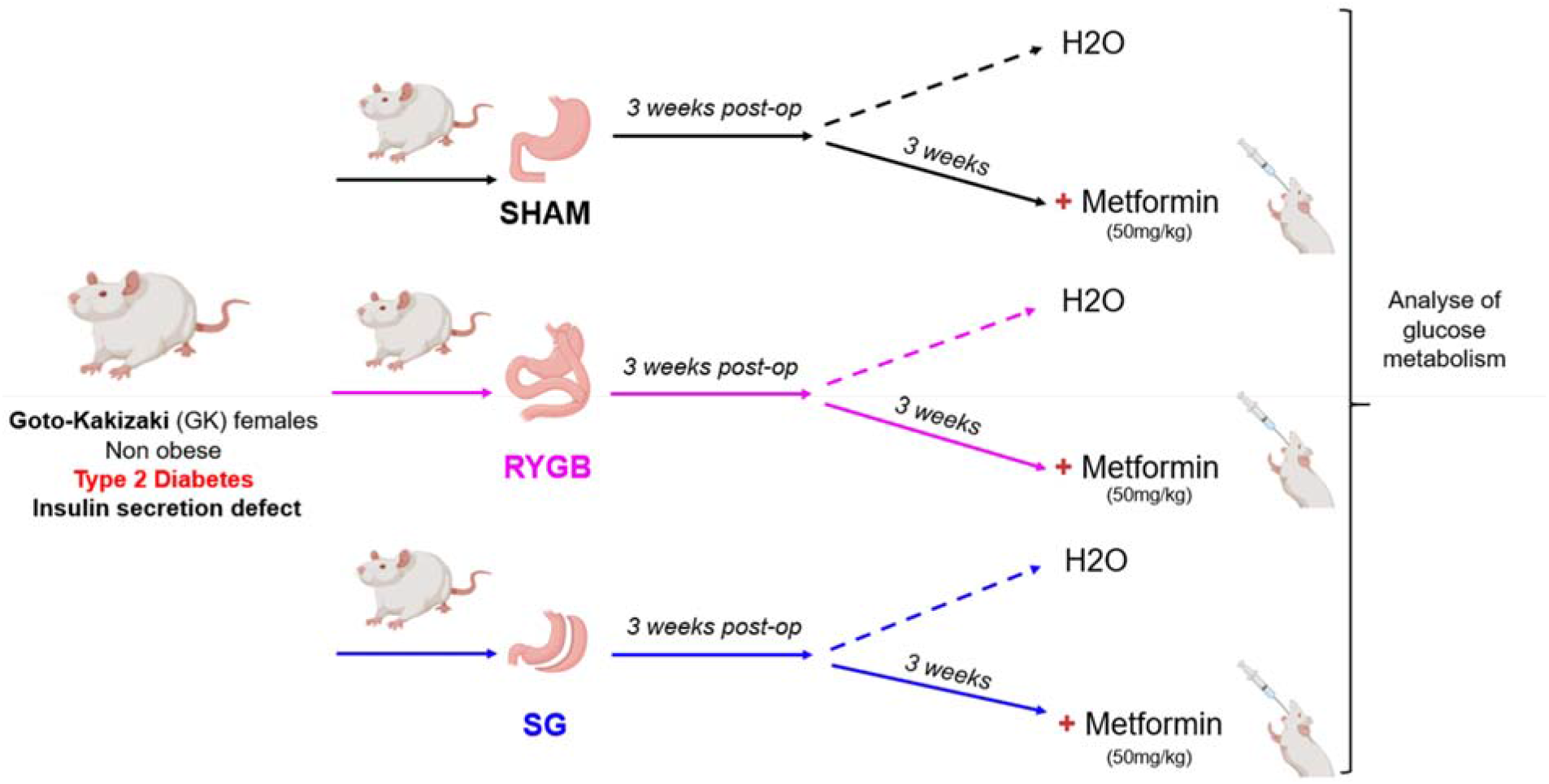
Study design. Schematic representation of the experimental timeline. Female GK rats underwent RYGB, SG, or sham surgery. Three weeks later, animals received metformin (50 mg/kg/day, 5 days/week) or vehicle for three additional weeks. Glucose tolerance and insulin sensitivity were assessed at week 6.

Metformin (Sigma-Aldrich, #317240-5GM) was administered by oral gavage at 50 mg/kg/day, 5 days/week, for 3 weeks. Control animals received water by gavage. At week 6 (i.e. after 3 weeks of metformin), animals underwent a standardized meal test and an insulin tolerance test (ITT).

### Surgical procedures and postoperative care

Rats were fasted overnight and anesthetized with isoflurane (5% induction, 2.5% maintenance; Vetflurane, Virbac). RYGB, SG, or sham surgeries were performed as previously described (16). Postoperatively, rats were housed individually and provided water 24 h post-surgery, then they were switched to a high-fat diet (HFD) (D12451 45kJ% fat, SSNIFF) to mimic relapse patterns of T2D observed in humans. Body weight and food intake were monitored daily for 2 weeks, then bi-weekly. After 2 weeks, rats were regrouped (4–5 per cage) according to surgical procedure to promote animal welfare.

### Meal test

Rats were fasted for 16 h with free access to water. They were then gavaged with a solution containing glucose (0.8 g/kg) and oil (2 ml/kg; Isio4 Lesieur®). In metformin-treated animals, 50 mg/kg metformin was added to the glucose/oil solution. Blood glucose was measured (Accu-Check®) at baseline and at 10, 20, 30, 60-, 90-, 120-, and 180-min. Area under the curve (AUC) was calculated (GraphPad Prism, GraphPad Software, San Diego, USA). Plasma samples were collected at baseline (T0) and 20 min post-gavage for hormone analyses.

### Insulin tolerance test (ITT)

ITT was performed one week before surgery and at week 6 (after 3 weeks of metformin treatment in the relevant groups). Rats were fasted for 5 h, then injected intraperitoneally with insulin (1 U/kg, NovoRapid® FlexPen®). Blood glucose was measured from tail blood at 0, 15, 30, 60, 90, 120, and 180 min.

### Blood collection and hormone assays

Blood was collected in lithium heparin tubes (Microvette® CB 300 LH) pre-treated with DPP-4 inhibitor (Roche). Samples (400 µl) were centrifuged at 1500 g for 20 min at 4°C, and plasma was stored at –80°C. Plasma GIP was measured by ELISA (Millipore® EZRMGIP-55K). GLP-1 and leptin were assayed using MSD® multiplex (U-PLEX Custom Metabolic Group 1, K153ACM-2). Insulin was quantified by ultrasensitive ELISA (Eurobio Scientific #80-INSRTU-E01).

### Statistical analysis

Data are presented as mean ± SEM. Between-group comparisons were made using Kruskal–Wallis with Dunn’s post hoc test, or Mann–Whitney for two-group comparisons. Repeated measures were analyzed by two-way ANOVA with Geisser–Greenhouse correction and Tukey’s multiple comparisons test. A p-value <0.05 was considered statistically significant.

## RESULTS

### 1) Baseline characteristics in female GK rats according to the condition

During the first three days post-surgery, all groups experienced weight loss due to the post-operative protocol (Fig. 2A). Sham-operated animals subsequently regained weight, exceeding baseline levels by day 14, whereas RYGB and SG groups maintained significantly lower body weights until the end of the experiment. Accordingly, plasma leptin levels, 6 weeks post-surgery, were significantly reduced in both RYGB and SG animals compared with sham controls (Fig. 2C). Cumulative food intake was also lower in operated rats than in sham controls (Fig. 2B).

**Figure 2.**
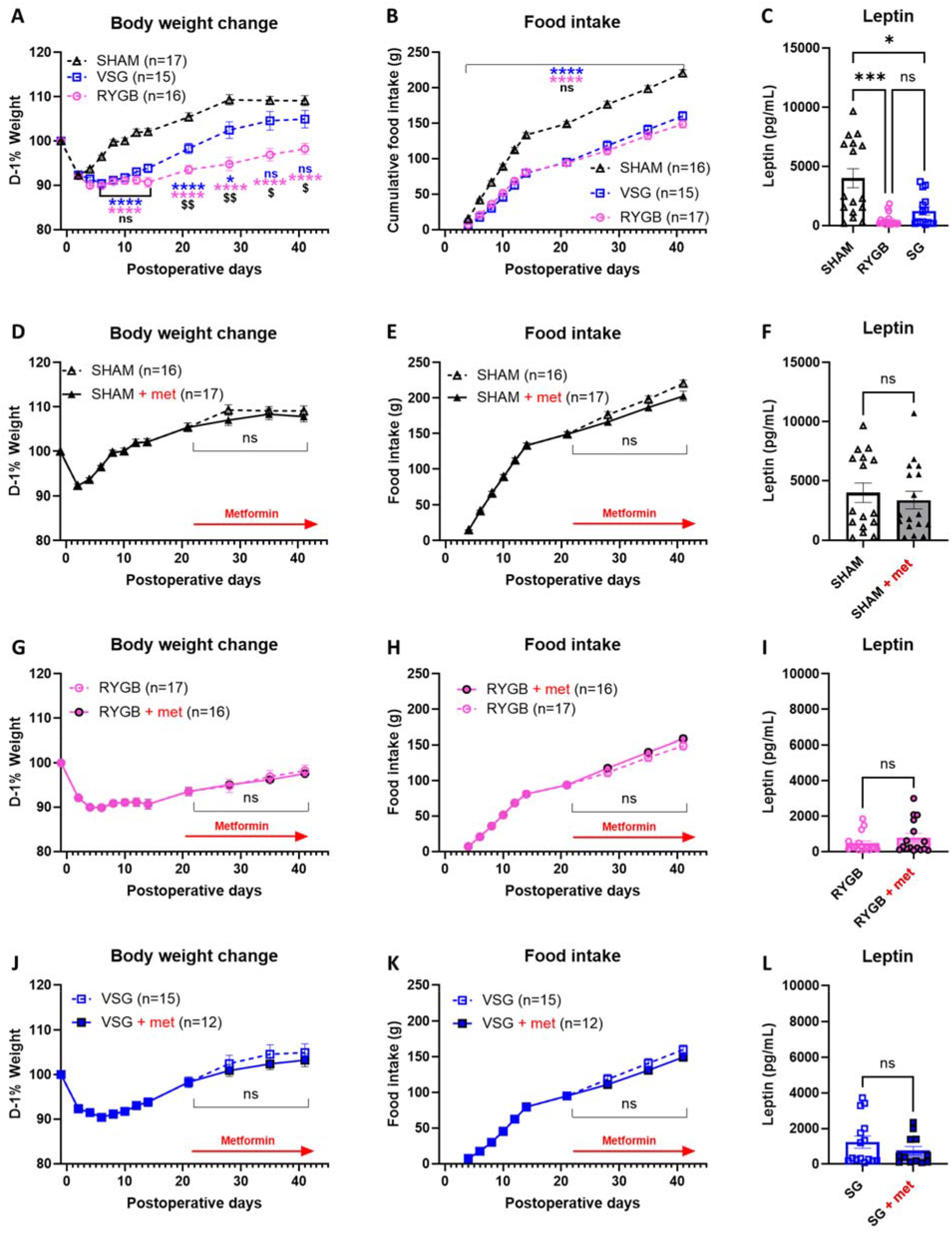
Effects of bariatric surgery and metformin on body weight, food intake, and leptin levels. (A–C) Effects of surgery alone: (A) Body weight change (% from baseline), (B) cumulative food intake, and (C) plasma leptin levels at day 40. (D–F) Effects of metformin in sham-operated rats. (G–I) Effects of metformin after RYGB. (J–L) Effects of metformin after SG. Data are mean ± SEM. SHAM n=16, RYGB n=17, SG n=15, SHAM+Met n=17, RYGB+Met n=16, SG+Met n=12. ^*^ p<0.05, ^***^ p<0.01, ^****^ p<0.001, ns no statistical difference

Metformin administration for three weeks, whether in sham-operated animals or following RYGB or SG did not further alter body weight trajectory, food intake, or plasma leptin compared with the respective surgery-only groups (Fig. 2D-L).

### 2) Bariatric surgery improves glucose tolerance, insulin sensitivity, incretin secretion, and insulin secretion

Fasting blood glucose levels were significantly lower in RYGB and SG rats compared to sham controls (Fig. 3A). Following oral glucose and lipid gavage, the amplitude of the glycemic peak was more pronounced in RYGB and SG operated rats compared to sham (Fig. 3B), although overall glycemic exposure evaluated by AUC did not differ significantly (Fig. 3C). Slope analyses revealed altered glycemic dynamics: both the ascending slope, reflecting blood glucose appearance, (Fig. 3D), and the descending slope, reflecting blood glucose clearance, were steeper (Fig. 3E) in operated rats compared to sham animals. Postprandial GIP and GLP-1 secretion increased significantly after both surgeries, whereas fasting levels remained unchanged (Fig. 3H–I). Post-prandial insulin concentrations slightly increased in SG compared with the Sham or RYGB (Fig. 3J).

**Figure 3.**
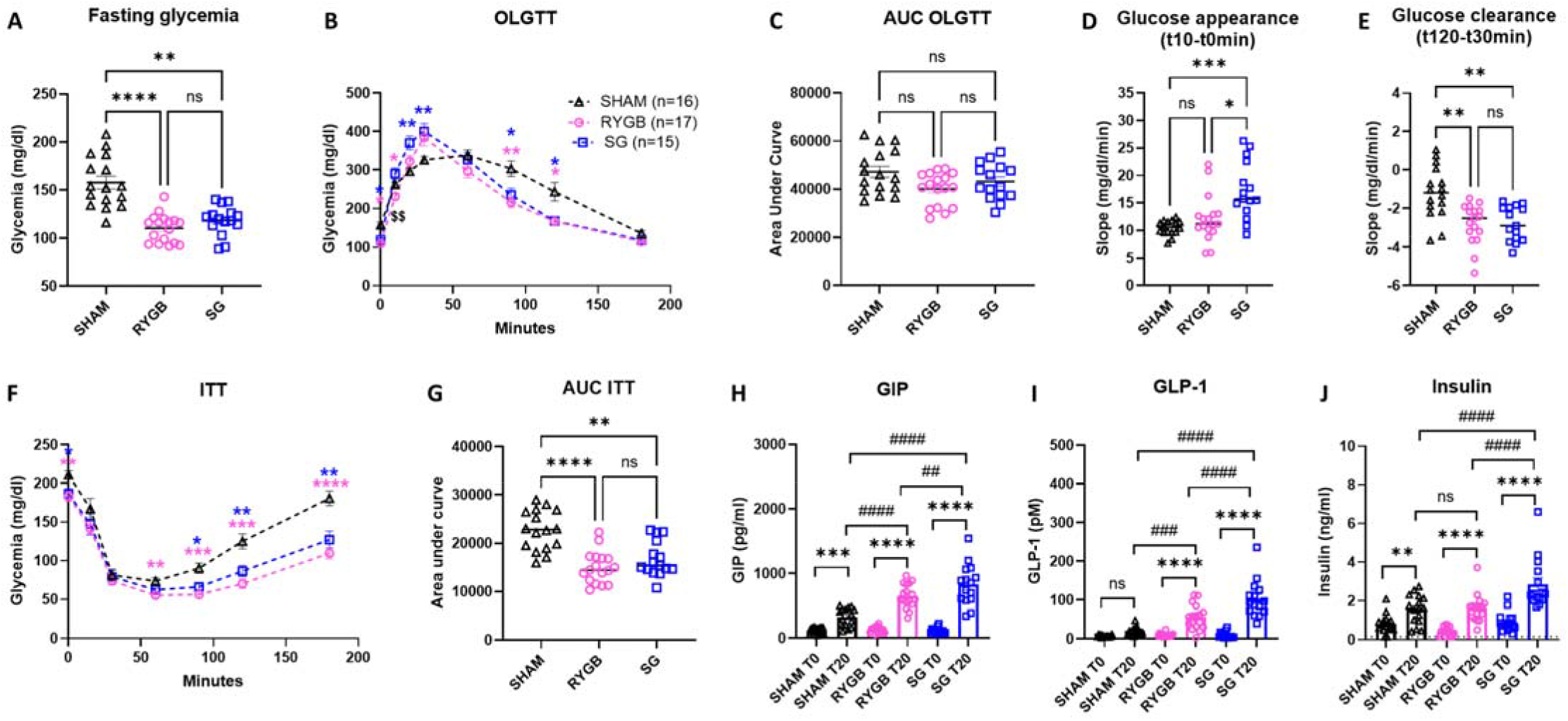
Bariatric surgery improves glucose homeostasis. (A) Fasting glucose. (B-C) Postprandial glucose curves (OGLTT) and corresponding Area under the curve (AUC). (D–E) Glucose appearance and clearance corresponding to the ascending and descending slopes of the glycemic curve. (F–G) Insulin tolerance test (ITT) and corresponding AUC. (H–J) Preprandial (T0) and postprandial (T20min) plasma levels of GIP, GLP-1, and insulin. SHAM n=16, RYGB n=17, SG n=15. ^*^ p<0.05, ^**^ p<0.01^***^ p<0.001, ^****^ p<0.0001 # p<0.05, ## p<0.01, ### p<0.001, #### p<0.0001 ns : no statistical difference

Insulin tolerance tests (ITT) revealed improved insulin sensitivity in RYGB and SG compared with sham, evidenced by a more pronounced blood glucose clearance and delayed recovery (Fig. 3F-G).

### 3) Metformin improves glucose tolerance and insulin sensitivity, but not incretin or insulin secretion

After three weeks of metformin treatment, fasting glucose levels showed a non-significant trend toward reduction in metformin-treated sham rats (Fig. 4A). However, metformin significantly improved glucose tolerance, lowering postprandial glycemic excursions and reducing AUC on a meal test (Fig. 4B-C). The glycemic peak at 30 min was attenuated, although the slopes of the glycemic curve remained unchanged (Fig. 4D-H). No significant differences were observed in GIP, GLP-1 nor insulin levels between groups (Fig. 4H-J). Metformin modestly but significantly improved insulin sensitivity (Fig. 4F-G).

**Figure 4.**
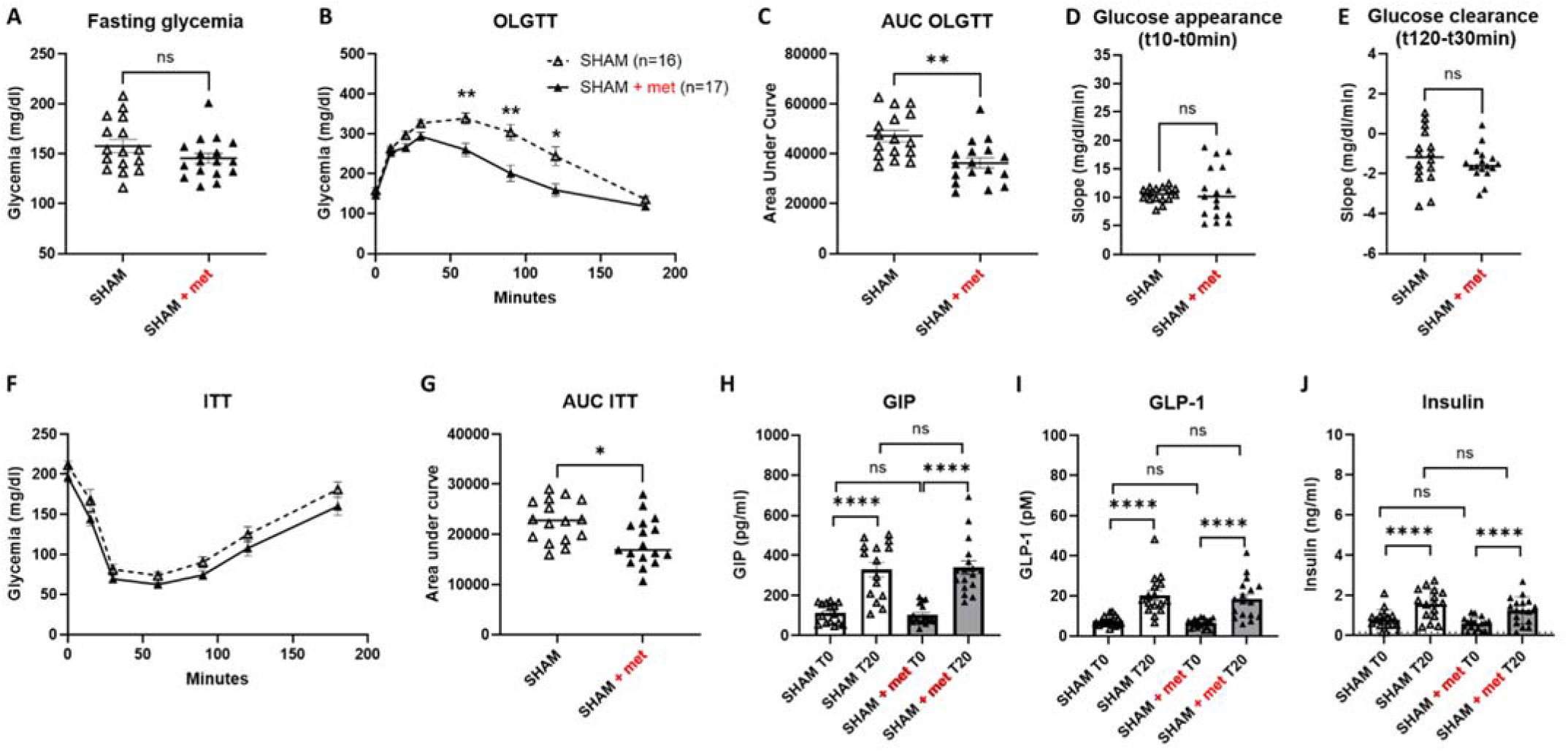
Metformin improves glucose tolerance and insulin sensitivity in sham-operated rats. (A) Fasting glucose. (B-C) Postprandial glucose curves (OGLTT) and corresponding Area under the curve (AUC). (D–E) Glucose appearance and clearance corresponding to the ascending and descending slopes of the glycemic curve. (F–G) Insulin tolerance test (ITT) and corresponding AUC. (H–J) Preprandial (T0) and postprandial (T20min) plasma levels of GIP, GLP-1, and insulin. SHAM n=16, SHAM+Met n=17. * p<0.05, ** p<0.01^***^ p<0.001, ^****^; p<0.0001 # p<0.05, ## p<0.01, ### p<0.001, #### p<0.0001 ns : no statistical difference

### 4) Combination of bariatric surgery and metformin improves glucose tolerance, but not insulin sensitivity, incretin, or insulin secretion

Addition of metformin did not affect fasting glucose levels after RYGB or SG (Fig. 5A-B 6A-B). However, glucose tolerance was significantly improved compared to surgery alone, as evidenced by reduced AUC and an earlier glycemic peak at 20 min during the meal test (Fig. 5B-C, 6B-C). The slopes of the glycemic curves weren’t different between surgery and surgery + metformin (Fig. 5D-E, 6D-E). Metformin did not further increase the surgery-induced postprandial GIP and GLP-1 secretion, nor did it alter insulin levels (Fig. 5H-J, 6H-J). ITT showed no additional improvement of metformin in insulin sensitivity beyond that achieved by surgery alone (Fig. 5F-G, 6F-G).

**Figure 5.**
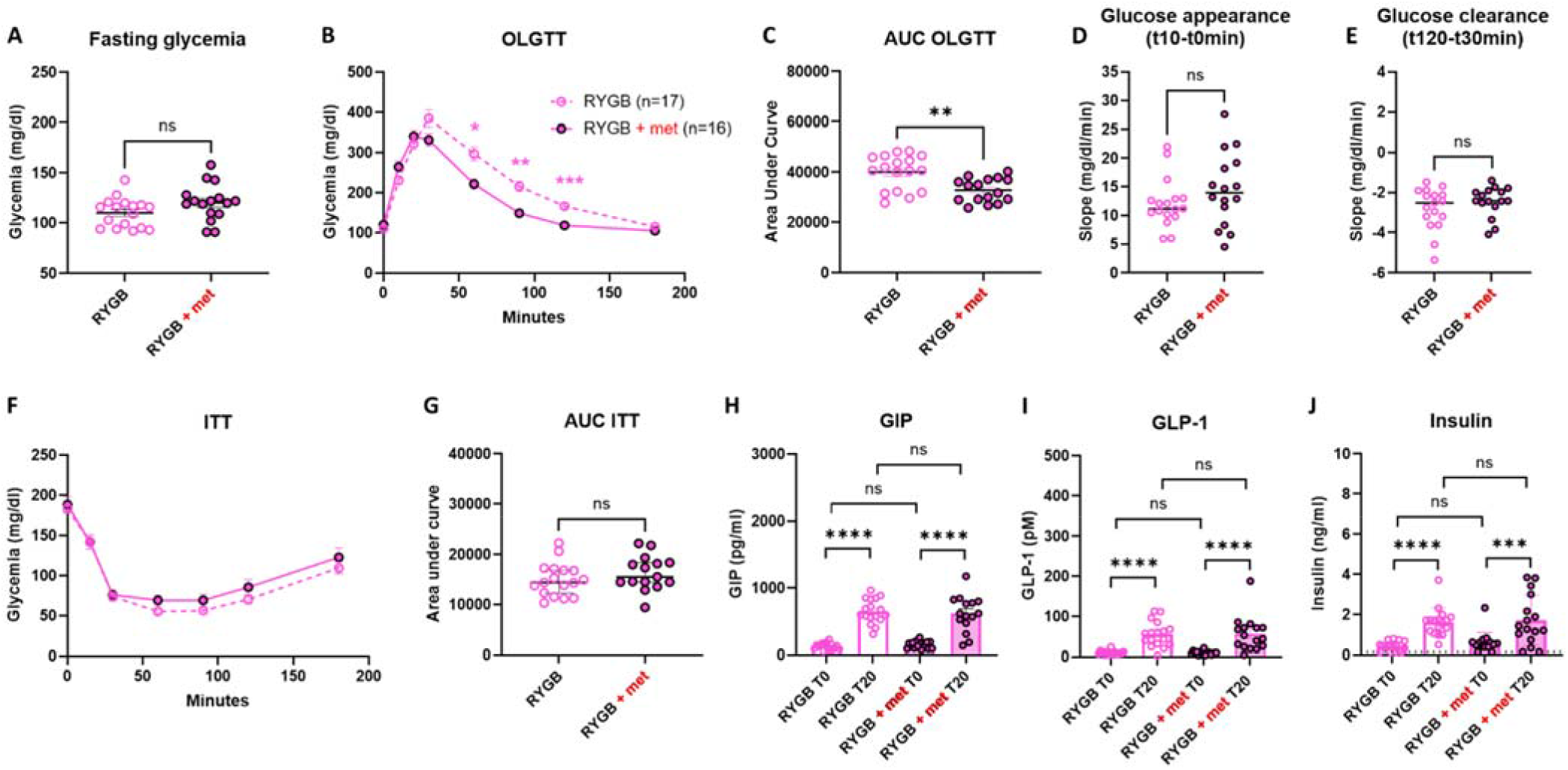
Combined effects of RYGB and metformin. (A) Fasting glucose. (B-C) Postprandial glucose curves (OGLTT) and corresponding Area under the curve (AUC). (D–E) Glucose appearance and clearance corresponding to the ascending and descending slopes of the glycemic curve. (F–G) Insulin tolerance test (ITT) and corresponding AUC. (H–J) Preprandial (T0) and postprandial (T20min) plasma levels of GIP, GLP-1, and insulin. ^*^ p<0.05, ^**^ p<0.01^***^ p<0.001, ^****^ p<0.0001 # p<0.05, ## p<0.01, ### p<0.001, #### p<0.0001 ns : no statistical difference

**Figure 6.**
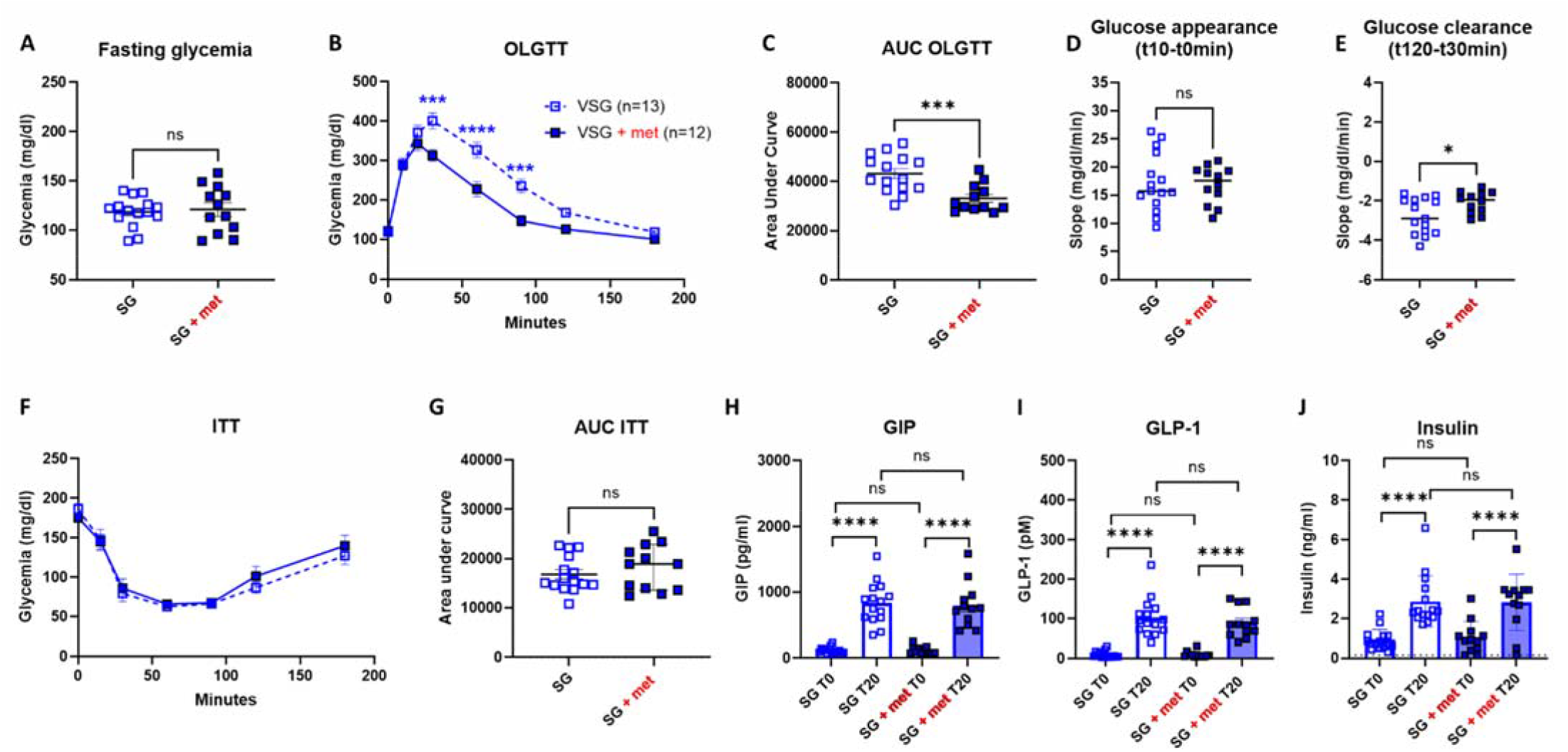
Combined effects of SG and metformin. (A) Fasting glucose. (B-C) Postprandial glucose curves (OGLTT) and corresponding Area under the curve (AUC). (D–E) Glucose appearance and clearance corresponding to the ascending and descending slopes of the glycemic curve. (F–G) Insulin tolerance test (ITT) and corresponding AUC. (H–J) Postprandial plasma levels of GIP, GLP-1, and insulin (T0 vs T20). SG n=15, SG + Met n=12 Blue asterisk, P<0.05 between SG and SG+Met

## DISCUSSION

In this study, we show that bariatric-metabolic surgery (BMS) significantly improves glucose homeostasis in a non-obese and insulin-deficient model of type 2 diabetes, the female Goto-Kakizaki (GK) rat. Both RYGB and SG led to reductions in body weight, food intake, and leptin levels, alongside improvements in fasting glycemia, glucose tolerance, and insulin sensitivity. These effects were accompanied by enhanced postprandial secretion of incretins GLP-1 and GIP, but occurred without complete restoration of insulin secretion, consistent with the intrinsic β-cell defect characteristic of the GK model. The accentuated postprandial glycemic peak and steeper glycemic slopes observed after surgery may reflect accelerated gastric emptying or enhanced intestinal glucose absorption, mechanisms previously described in both rodents and humans following BMS (17). However, as neither gastric emptying nor intestinal glucose uptake were directly assessed in this study, this interpretation remains speculative.

Interestingly, SG induced greater postprandial incretin and insulin secretions than RYGB in our model which contrasts with other preclinical studies (18) and clinical data where RYGB typically elicits stronger hormonal responses (6). This discrepancy may be attributed to specie-specific responses, the non-obese diabetic phenotype of GK rats, or sex-related metabolic variations, as our study focused exclusively on females, a population underrepresented in preclinical metabolic research. This sex-specific design is clinically relevant, as women represent the majority of patients undergoing bariatric surgery and show differences in response to surgery (19, 20). Moreover, sexual dimorphism plays a significant role in metabolic regulation, influencing adipose tissue distribution, insulin sensitivity, and incretin responses (13, 21). Future studies directly comparing male and female GK rats will be essential to delineate the contribution of sex hormones and genetic factors to postoperative metabolic outcomes.

Metformin treatment in sham-operated animals improved glucose tolerance and insulin sensitivity, consistent with its known mechanisms of action particularly the reduction of hepatic gluconeogenesis (22). Notably, these effects were achieved using a relatively low dose (50 mg/kg), chosen to approximate human therapeutic exposures and avoid the supra-pharmacological doses often used in rodent studies (23). In contrast to some reports, metformin did not increase GLP-1 secretion in our study. This may be due to the early postprandial sampling time (20 min), whereas metformin-induced GLP-1 elevations are typically observed later (around 60 min) (24). When combined with surgery, metformin further improved postprandial glucose handling by reducing glycemic excursions and shifting the timing of the glycemic peak to an earlier time point. However, no additive effects were observed on insulin sensitivity, incretin secretion, or insulin release, suggesting that these mechanisms may already be maximally engaged by surgery alone. Thus, the primary benefit of adding metformin in this context appears to be the attenuation of postprandial glucose excursion. Importantly, inter-individual variability in response to combined treatment suggests the existence of responder and non-responder profiles, warranting further investigation.

The strengths of our study include the use of a non-obese, insulin-deficient model of T2D that mirrors key features of the SIDD cluster in humans and allows us to disentangle diabetes from obesity. Furthermore, by focusing on female rats, we addressed a gap in the literature, as most previous preclinical studies have relied on males, despite growing evidence of sex-specific metabolic responses (25) (26) (27). However, several limitations should be acknowledged. First, the study duration was relatively short (six weeks), which may not be sufficient to capture delayed effects of surgery on insulin secretion (25). Accordingly, previous studies in GK male rats suggest that longer follow-up may be required to detect changes in insulin secretion after surgery (25) (26) (27).

In conclusion, our findings demonstrate that both bariatric surgery and metformin independently improve glucose regulation in a non-obese, insulin-deficient model of type 2 diabetes. While surgery alone maximizes improvements in insulin sensitivity and incretin secretion, the addition of metformin provides further benefit by attenuating postprandial glycemic excursions and advancing the timing of the glycemic peak. These effects may help reduce the risk of postprandial hyperglycemia and subsequent hypoglycemic episodes, which are complicated problems to deal with after BMS even if it hasn’t yet been tested on humans (28).

Importantly, our study highlights the potential of metformin as a valuable adjunct to bariatric surgery in non-obese, insulin-deficient diabetes. The use of female GK rats addresses a critical gap in preclinical research and aligns with the clinical reality that women represent most bariatric surgery recipients. Future studies should extend the duration of the follow-up, to assess long-term effects on β-cell function and glycemic control include both sexes to explore sex-specific responses and investigate individual variability to identify responder profiles. These steps will be essential to enhance translational relevance of our finding for SIDD patient population.

## DATA AVAILABILITY

The source data are available to verified researchers upon request by contacting the corresponding author.

## ACKNOWLEDGMENTS

The authors thank the staff of CRI and BFA animal facilities for their expert care and support throughout the study. We thank André Bado, Johanne Le Bihan Le Beyec, and Séverine Ledoux for their important inputs and discussions.

## AUTHOR CONTRIBUTIONS

Mélanie Hirlemann: Conceptualization, Investigation, Writing - Original Draft, Mélanie Garmon: Investigation, Lara Ribeiro-Parenti: Methodology, Danielle Bailbé: Resources, Methodology, Alexandra Willemetz: Investigation, Hounayda El Jindi Shahrour: Investigation, Jamileh Movassat: Resources, Methodology, Claire Carette: Conceptualization, Project administration, Funding acquisition, Writing - Review & Editing, Maude Le Gall: Conceptualization Project administration Funding acquisition Writing - Review & editing

